# A modular microreservoir for active implantable drug delivery

**DOI:** 10.1101/762716

**Authors:** Farzad Forouzandeh, Xiaoxia Zhu, Nuzhet N. Ahamed, Joseph P. Walton, Robert D. Frisina, David A. Borkholder

## Abstract

Active implantable microscale reservoir-based drug delivery systems enabled novel and effective drug delivery concepts for both systemic and localized drug delivery applications. These systems typically consist of a drug reservoir and an active pumping mechanism for precise delivery of drugs. Here we present a stand-alone, refillable, scalable, and fully implantable microreservoir platform to be integrated with micropumps as a storing component of active implantable drug delivery microsystems. The microreservoir was fabricated with 3D-printing technology, enabling miniature, scalable, and planar structure, optimized for subcutaneous implantation especially in small animals (e.g., mouse), while being readily scalable for larger animals and human translation. Three different capacities of the microreservoir (1 μL, 10 μL, and 100 μL) were fabricated and characterized all with 3 mm thickness. The microreservoir consists of two main parts: a cavity for long-term drug storage with an outlet microtubing (250 μm OD, 125 μm ID), and a refill port for transcutaneous refills through a septum. The cavity membrane is fabricated with thin Parylene-C layers using a polyethylene glycol sacrificial layer, minimizing restoring force and hence backflow, as fluid is discharged. This feature enables integration to normally-open mechanisms and improves pumping efficiency when integrated to normally-closed pumps. The results of *in vitro* optimization and characterization of the cavity membrane show 95% extraction percentage of the cavity with insignificant (2%) backflow due to restoring force of the membrane. The refill port septum thickness is minimized down to 1 mm by a novel pre-compression concept, while capable of ~65000 injections with 30 Ga non-coring needles without leakage under 100 kPa (4× greater than physiological backpressure). To demonstrate integrability of the microreservoir to an active micropump, the 10 μL microreservoir was integrated to a micropump recently developed in our laboratory, making an implantable drug delivery microsystem. Two different microsystems were subcutaneously implanted in two mice, and the outlet microtubing was implanted into the round window membrane niche for infusion of a known ototoxic compound (sodium salicylate) at 50 nL/min for 20 min. Real-time shifts in distortion product otoacoustic emission thresholds and amplitudes were measured during the infusion. The *in vivo* results show a mean shift of 22.1 dB after 20 min for the most basal region, matching with syringe pump results. A biocompatibility experiment was performed on the microsystem for six months to assess design and fabrication suitability for chronic subcutaneous implantation and clinical translational development. The results demonstrate very favorable signs of biocompatibility for long-term implantation. Although tested here on mice for a specific inner ear application, this low-cost design and fabrication methodology is scalable for use in larger animals and human for different applications/delivery sites.

## 1. Introduction

Drug delivery systems have become of interest to researchers in the past decades for improving therapeutic response by providing more consistent blood level compared to immediate release or sustained release parenteral depot, enabling less frequent and more efficient dosing, and improving patient comfort, safety, and compliance. Microscale reservoir-based delivery systems result in miniaturization of the system while allowing more precise control of the delivery rate for both systemic and site-directed delivery [1]. Specifically, these systems assist site-directed delivery by enabling access to relatively inaccessible sites or a specific tissue type, along with limiting side effects of systemic exposure. Microscale reservoir-based delivery systems can be categorized into three types: oral, dermal, and implantable delivery. This work focuses on the latter.

Implantable microscale reservoir-based drug delivery systems enabled novel and effective drug delivery concepts for both systemic and targeted drug delivery applications[1]. Implantable drug delivery systems have been classified into two categories [2–6]: (i) passive, where post-implantation drug release control is not feasible since it is pre-determined by the fabrication methods, materials, and drug formulation, and (ii) active, where post-implantation drug release control is enabled using electrical, mechanical, magnetic or other actuation methods. Passive implantable drug delivery systems are driven by osmotic potential or diffusion. These microsystems have been employed in various applications such as ophthalmology [7–13], oncology [14–16], interventional cardiology [17–20], precocious puberty [7], acromegaly[21], Hepatitis C [21–23], prostate cancer [24–26], pain indications [27], type 2 diabetes and Hepatitis [28–32]. In contrast, active implantable drug delivery systems are driven by mechanical pumping [33,34], electrolysis[35], and other actuation methods, which enable the patient or physician to start/modify/stop drug release interactively. These systems have been used for emergency care [36,37], treatment of ocular diseases [38,39] and cancer [40], and general application in small animals [41]. Active implantable drug delivery systems generally consist of a drug reservoir and a micropump for precise delivery of small volumes of a drug from the reservoir to the target organ. These systems are often normally-closed or have a check valve in the reservoir downstream.

The drug reservoir is an often overlooked component of active implantable drug delivery microsystems. Depending on the application, drug reservoirs can be refillable or non-refillable. Non-refillable microsystems are usually disposable; they are filled with the drug, implanted, and actuated to discharge the drug. These microsystems have been used in insulin delivery [42,43], ophthalmology [44], and emergency care [36,37]. In most of the active implantable drug delivery systems, the drug reservoir is refillable, enabling physicians to transcutaneously refill the reservoir via injections through sharp, thin, non-coring needles into septa in a designated refill port in the system housing [45,46]. The septa are usually made of silicone rubber[45] with self-healing[47] capabilities due to high resilience[48] and high deformability and self-adherence[49,50]. The ports typically have a base plate for penetration depth limitation and a cavity between the septum and base plate with an exit channel to provide a fluidic connection to the cavity membrane[51–53]. In some cases, a raised ridge[53,54] or a ring with different color[39] is incorporated to assist palpation or visual identification for the surgeon.

Although there are several microsystems in the literature or commercially available that use a septum for refilling [53,55], only a few of them report the number of punctures they can withstand without leakage. The first refillable microelectromechanical (MEMS) drug delivery device was developed by Lo et al. for the treatment of ocular diseases [38,39,56,57]. Refilling could take place through the top surface of the microsystem, made of polydimethylsiloxane (PDMS), while a Polyether ether ketone baseplate was embedded to control the penetration depth. The refilling port prevented leakage until up to 24 punctures under ~5 and ~30 kPa backpressures for membrane thicknesses of 250 μm and 673 μm, respectively. PROMETRA® programmable pumps (InSet Technologies Incorporation, NJ, USA) use silicone rubber for the septum, rated for withstanding an average of 1000 punctures. Hamilton® GC Septa are rated for a maximum of 100 injections using a 26s Ga needle, with a minimum thickness and diameter of 3 and 5 mm.

The dynamics between the fluid stored in the reservoir and the downstream environment when the pump is deactivated is an important feature in the design of the reservoirs. Since the reservoir membrane material is typically made of rubber-like or metal materials, extraction of the drug causes a restoring force. This restoring force can result in a potential backflow, which in the literature has been compensated by either external compression of the membrane [58] or replacement of the drug with electrolysis gases [57]. If not compensated in the cavity area, the backflow is usually prevented with normally-closed micropumps between the reservoir and the downstream system [53], or check valves [39,41]. To the best of our knowledge, there is no reported microsystem utilizing a cavity membrane with negligible restoring force to integrate to micropumps.

Laboratory animal models play an important role in understanding and treating human diseases. The most common animal models (approximately 95%) in research are rats and mice[59]. However, mice are preferred since they are easier to be genetically manipulated [60]. Further, for subcutaneous implantation to be a minor procedure, the surgery should involve less than 10% of the animal’s surface area [61]. These considerations necessitate the need for both miniaturization and scalability of a microsystem for experiments in small animal models (e.g., mice), and later translation to humans. Most of the reported studies and commercially available drug delivery microsystems have the reservoir and pumping mechanism integrated [41,55]. Due to limitations for scalability of pumping mechanism, especially for MEMS-based devices, scalability of these devices is challenging. This necessitates the development of a stand-alone implantable microreservoir with different capacities for integration to different micropumps with different flow rates.

Here we present a stand-alone, refillable, scalable, and fully implantable microreservoir platform to be integrated with micropumps as the storing component of an active implantable drug delivery microsystem. This microreservoir platform is intended to have a small footprint and a planar form factor for subcutaneous implantation, especially in small animals (e.g., mice), while being readily scalable for larger animals and human translation. Using 3D-printing technology to fabricate the microreservoir structure, three different capacities of the microreservoir are demonstrated and characterized (1 μL, 10 μL, and 100 μL). The cavity membrane is fabricated with thin Parylene-C layers using polyethylene glycol (PEG) sacrificial layer and optimized for minimal restoring force as fluid is discharged. The septum thickness is minimized by a novel pre-compression concept enabling thinning of the overall system, while capable of thousands of refills without leakage. To demonstrate integrability of the microreservoir design, it was integrated into a micropump reported in our previous work [62] and implanted in mouse models for inner ear drug delivery. Results indicate functional round window membrane (RWM) drug delivery. Long-term biocompatibility was assessed *in vivo* over six months.

## 2. Materials and methods

### 2.1. System operation concept and architecture

To build a microscale, yet scalable structure, stereolithography (SLA) fabrication approach was employed. Formlabs SLA 3D-printer (Form 2, Formlabs Inc., MA, USA) was used to fabricate a rigid biocompatible structure comprising a refill port, cavity area, an outlet port, microchannels to connect them, and a baseplate. This rigid baseplate controls the penetration depth of the refill needle while provides a surface to form the cavity membrane. The rigid frame protects the enclosure and ensures sufficient rigidity for handling.

To provide a completely inert and biocompatible environment for therapeutic compounds, the stored drug was designed to be encapsulated with parylene, which is a USP Class VI material suitable for implant construction. Furthermore, parylene has less permeability to liquid compared to other common materials used in the literature for reservoir membrane, such as PDMS [38] and MDX-4-4210 [41]. After the first layer of parylene deposition, Molten PEG was deposited on the cavity area and solidified to build a biocompatible sacrificial layer, a technique that has been used to form parylene-made cavities [41]. A second parylene layer was deposited on the PEG sacrificial layer to build a thin pre-formed dome-shape cavity membrane. The pre-formed shape and thinness of the cavity membrane enable discharge of the fluid and collapse of the cavity membrane without having a significant restoring force that would result in original shape recovery and undesirable backflow from the outlet. The seal between two parylene layers is enhanced using a compressed biocompatible silicone-made gasket on the cavity edge. The thicknesses of the cavity membranes for three different capacities (1, 10, and 100 μL) were optimized for 100 kPa backpressure, which is four times larger than maximum physiological backpressure in human [63]. This backpressure was used as a criterion for both cavity membrane and septum characterization. For all the cavity sizes, the overall thickness of the microreservoir is set to 3 mm, which was shown to be suitable for long-term subcutaneous implantation in mice. An overview of the microreservoir concept is illustrated in Figure 1.

**Figure 1.**
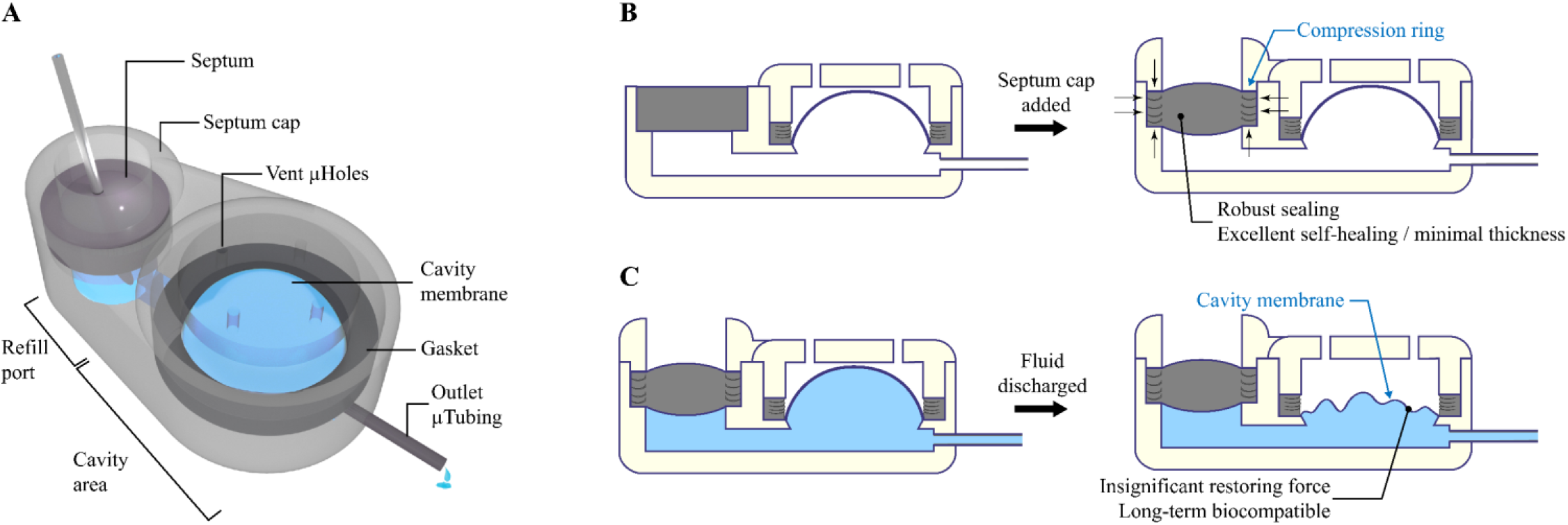
A) Conceptual schematic of the microreservoir. The structure of the microreservoir is made of two main sections: the refill port for refilling the microreservoir by injections through a septum, and the cavity area for storing the drug. The structure is made with 3D-printing technology using a biocompatible resin, enabling printing micro-scale features that are readily scalable, enabling applications from mice to humans. B) The septum cap incorporates a compression ring that causes the septum to be compressed both vertically and laterally. The vertical compression ensures sealing from the bottom and holds the septum during multiple injections. The lateral compression induces internal stress in the septum to improve its performance with numerous injections without leakage. This feature enables reduction of the septum thickness while maintaining high resistance to numerous punctures. C) The cavity membrane is made of a pre-formed dome-shape thin parylene layer, enabling negligible restoring force after partial/full discharge of the cavity when the pumping mechanism is switched off. This feature facilitates the integration of the microreservoir to normally-open micropumps and eliminates the necessity of using check valves for low-backpressure environments. Further, when integrated to normally-close micropumps, the constant backpressure on the check valves are eliminated and the pumping efficiency increases. Finally, since the entire internal surface of the microreservoir, including the septum and the cavity membrane, is coated with a parylene layer, long-term biocompatibility for drug storing is ensured.

The refill port has a 2.5 mm hole for the septum and a built-in septum stopper to ensure space between the septum and the baseplate, allowing fluid flow to the cavity area. The 2.5-mm septum is micro-molded in cylindrical shape using a biocompatible silicone rubber followed by parylene deposition, to provide complete encapsulation of the drug by the parylene. A cap with a raised compression ring was placed and fixed on the septum to provide vertical compression which improves sealing and facilitates fixation of the septum in the port during injection, which was a challenge in previous reports [64,65]. The compression cap also causes the septum to laterally push the port wall, inducing lateral compression in the septum, enhancing its self-healing and therefore, increasing its endurance to punctures without leakage. This feature facilitates a significant reduction of the septum thickness, enabling septum down to 1mm thickness to be successfully used. A raised ridge is designed on top of the septum cap to enable subcutaneous palpation and visual access over shaved skin, for performing refills.

### 2.2. Fabrication process

The microreservoir substrate was 3D-printed (Form 2, Formlabs Inc., MA, USA) using a rigid biocompatible material (Dental SG resin, Formlabs Inc., MA, USA) creating a hole of 2.5-mm diameter and 1-mm depth for the septum, cavity area, outlet port, fluidic interconnections between them, and a baseplate. In the cavity area, a raised circle-shaped ring was considered for placing the gasket, with an angled wall to the baseplate to reduce the cavity dead volume. In the refill port, a septum stopper with a diameter of 1.8 mm was designed, providing a seat for the septum. A polyurethane-based catheter microtubing (ID=125 μm, OD=250 μm; Micro-Renathane® Catheter Tubing, Braintree Scientific Inc., MA, USA) was fixed and sealed to the output port, using cyanoacrylate. The structure was coated with a 1-μm-thick layer of parylene using a PDS 2010 LABCOATER™ 2 (Specialty Coating Systems, Indianapolis, IN, USA).

Cavities with three different capacities (1, 10, and 100 μL) were fabricated, where the diameters of the cavity area for each of them were empirically found to be 1.3, 3.1, and 10 mm, considering 3 mm objective for the overall thickness. To build the cavity membrane with parylene-C, a sacrificial layer is required, which defines the capacity of the cavity after release. PEG (1,500 Mn, melting point: 37 °C, Sigma-Aldrich, MO, USA) was chosen for the sacrificial layer due to its biocompatibility and solubility in water. PEG was melted at 70 °C on a hotplate and mixed with a food dye (McCormick Food Color and Egg Dye, McCormick & CO., MD, USA) for visual confirmation of its release. Using a heated micropipette (70 °C), PEG was deposited on the cavity area which was held at room temperature and treated with hydrophobic spray (Scotchgard™ Fabric & Upholstery Protector, 3M Co, MN, USA) for quick solidification of PEG and avoiding its flow into the space under the septum and the gasket raised ring. It was empirically found that keeping the substrate at room temperature can result in smooth PEG surfaces, while providing a sufficiently fast cooling rate to avoid the flow of the PEG in the channel regions. The full volume of required PEG for different capacities were deposited with a single micropipette dispense.

To make the cavity membrane on top of the PEG sacrificial layer, the required thickness of the second parylene layer for different capacities was estimated based on thin-walled spherical pressure:

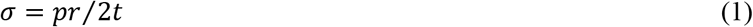

where σ, p, r, and t represent tensile stress on the membrane, internal pressure, cavity radius, and membrane thickness, respectively. The membrane is designed for 100 kPa backpressure (p= 100 kPa), while the radius was determined based on overall thickness and capacity of the device. The thicknesses of the parylene membrane for each cavity size was calculated to achieve membrane stresses smaller than the tensile strength of parylene (69 MPa [66]) by a factor of 5. The thicknesses of deposited parylene membranes were 2.7, 5.6, and 18.1 μm for 1, 10, and 100 μL capacities respectively.

Silicone rubber (MED-6215, NuSil™ Technology LLC, CA, USA) gaskets with 0.5 mm width and height with inner diameters of 1.3, 3.1, and 10 mm for capacities of 1, 10, and 100 μL, respectively, were micro-molded using a parylene-coated aluminum mold fabricated with conventional machining. The gaskets were cured at 150 °C for 15 minutes. The gasket was placed on the cavity area and was fixed and compressed using a parylene coated cavity cap which was 3D-printed (Form 2, Formlabs Inc., MA, USA). The cavity cap also covers the cavity membrane to protect it from external mechanical stress. The cavity cap included vent holes with 0.2-mm diameter to allow ingress and egress of air or liquid from the space between the cap and the cavity membrane during filling and discharging. These holes are smaller than the smallest needle size for this microreservoir (30 Ga, 311 um OD) to avoid inadvertent puncture of the membrane during subcutaneously refilling. The use of class VI biocompatible silicone rubber for the gasket and parylene deposition of the cavity caps minimizes the risk of inflammation with direct contact to body tissues.

Under visual observation using a microscope (Motic SMZ-168), the substrate was heated on a hotplate at 70 °C to melt the PEG, allowing it to be rinsed away with 100 mL 70°C deionized (DI) water. A silicone septum with 2.5 mm diameter and 1 mm thickness was micro-molded using parylene-coated 3D-printed molds and cured at 150 °C for 15 minutes and were coated with 1 μm parylene layer. The septum was placed in the refill port followed by placing and affixing a 3D-printed septum cap using cyanoacrylate. The septum cap incorporates a 0.2-mm-thick compression ring (1.8 mm ID, 2.5 mm OD) to compress the septum to provide internal stress for improving its self-healing properties. A raised ridge (1.8 mm ID, 3 mm OD, 0.5 mm thickness) was integrated on top of the septum cap to facilitate palpation for subcutaneous refilling. All the surfaces were filleted to minimize potential mechanical inflammation after implantation. Dimension details of the 3D-printed structure are provided in Figure S1. The full fabrication process is illustrated in Figure 2.

**Figure 2.**
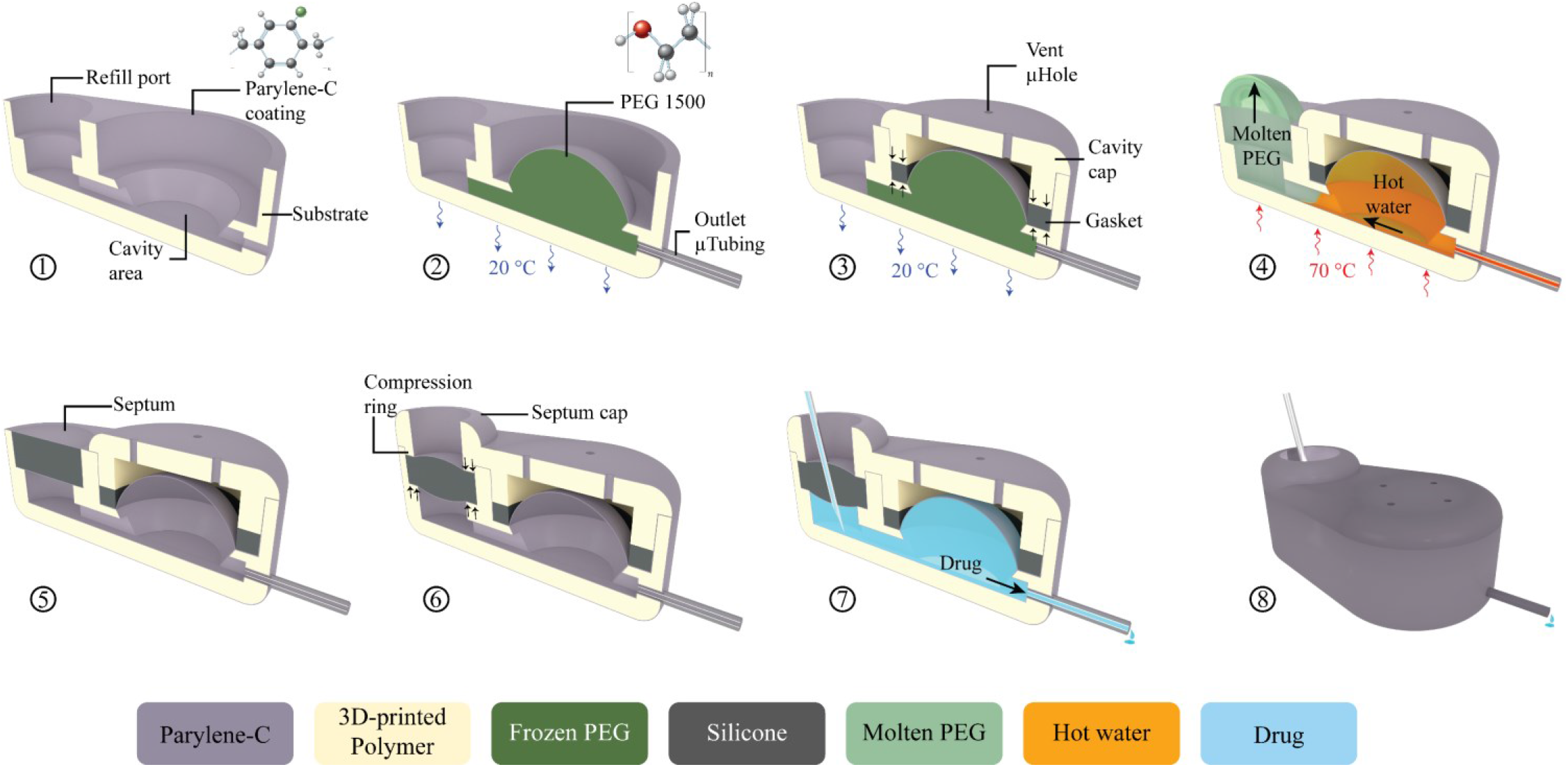
A cut-view schematic of the microreservoir fabrication process. 1) The substrate of the microreservoir was 3D-printed using a biocompatible resin, followed by 1 μm parylene-C deposition. 2)A microtubing (250 μm OD, 125 μm ID) was inserted into the substrate outlet port and sealed in place with cyanoacrylate. The substrate was placed on a cold plate at room temperature and molten PEG at 70 °C was deposited on the cavity area to solidify and create the microreservoir volume. This was followed by another Parylene-C deposition to create the deformable microreservoir membrane capable of withstanding 100 kPa backpressure. 3) A silicone gasket was fabricated using a micro-molding technique and was placed within the cavity around the PEG dome. A 3D-printed Parylene-C coated cap was affixed on top of the cavity area with cyanoacrylate, compressing the O-ring to reinforce sealing between the two Parylene-C layers, and to protect the cavity from mechanical stress. The cap has vents to allow egress/ingress of air or fluid between the membrane and the cap during filling/discharging. 4) The substrate was placed on a hotplate at 70 °C to melt the PEG and wash it by gentle injection of 100 mL of DI water at 70 °C 5) A 2.5-mm diameter, 1-mm-thick septum made of long-term implantable silicone rubber was micro-molded and coated with parylene-C. The septum was placed in the refilling port (2.5 mm diameter). 6) A 3D-printed, Parylene-C coated septum cap with an extruded compression (2.5 mm OD, 1.8 mm ID) was affixed on top of the septum with cyanoacrylate to compress the septum providing a sealing force on the bottom and sides while enhancing the self-healing properties when punctured with refilling needles. 7) schematic cut view of the fabricated microreservoir being filled by a refilling needle. 8) A schematic full-body view of the microreservoir.

To demonstrate integrability of the microreservoir to a micropump, a 10 μL microreservoir was integrated with an implantable micropump for inner ear drug delivery, recently developed in our group [62]. A miniature container was 3D-printed for the micropump right beside the microreservoir, while the overall thickness was kept at 3 mm. Fillets with 1 mm radius were applied to the container walls to provide curved surfaces to improve long-term subcutaneous implantability. The microreservoir was built following the fabrication process outlined in Figure 2. The micropump was placed in the container, and its microtubing was placed and sealed to the microreservoir outlet using cyanoacrylate. The micropump was then fixed in the container using cyanoacrylate, and the silicone rubber (MED-6215, NuSil™ Technology LLC, CA, USA) was poured in the container and cured at room temperature for 48 hrs. To soften the device surface, the entire microsystem except the cavity cap was covered with a thin layer of silicone by manually applying droplets of silicone to evenly coat the outer surface, and cured at room temperature for 48 hrs.

### 2.3. Benchtop experimental methods

To assess the functionality of the septum and the cavity membrane, these two parts were decoupled and tested separately. Two separate sets of test rigs were designed and fabricated and tested with samples fabricated following the fabrication process described in section 2.2.

#### 2.3.1. Septum test

The septum was tested utilizing separately fabricated septum samples, consisting of a 3D-printed refill port with septum stopper and space beneath it along with the outlet coupled to a Tygon tubing (0.508 mm ID, 1.52 mm OD). The septum and septum cap were then placed and fixed in the refill port. Although in some studies the septum leakage test is performed using N2 as the working fluid [64], here we tested the septum using dyed DI water since it simulates the *in situ* condition where the septum is in contact with a liquid drug.

A pneumatic puncture device was designed and fabricated to hold the septum sample and automatically puncture through the septum in a single location, to test the worst-case scenario [39,64]. The structure of the pneumatic puncture device was 3D-printed using Grey Pro resin (Formlabs Inc., MA, USA), having a groove for the test sample to be placed and fixed with a set screw beneath the pneumatic puncture mechanism. A miniature cylinder (SM-3-1, Clippard Co., OH, USA) was placed and fixed on top of the sample. An electronic pressure regulator (ITV2030-31N2BL4, SMC Co., Tokyo, Japan) was connected to the cylinder which was fed with sine waves of 0.2 Hz frequency by a signal generator, to simulate a realistic manual puncture speed. A needle was affixed to a 3D-printed needle holder which was press-fit to the piston of the cylinder.

Commercially available 27 and 30 Ga (413 and 311 μm OD) needles were used for the puncture tests. These sizes were selected as a trade-off between maximizing needle and septum lifetime: the needles need to be large enough to puncture through the skin and the septum without bending the tip, while small enough to minimize septum damage. Furthermore, the opening diameter of the septum cap is 1.8 mm, making needle sizes larger than 27 Ga impractical. The needles were machined to non-coring shape to minimize damage to the septum structure [38,39,56]. The needle tip was beveled to a point with a 12° angle recommended for animal injections [67]. Figure 3A shows the pneumatic puncture device with a 30 Ga needle.

**Figure 3.**
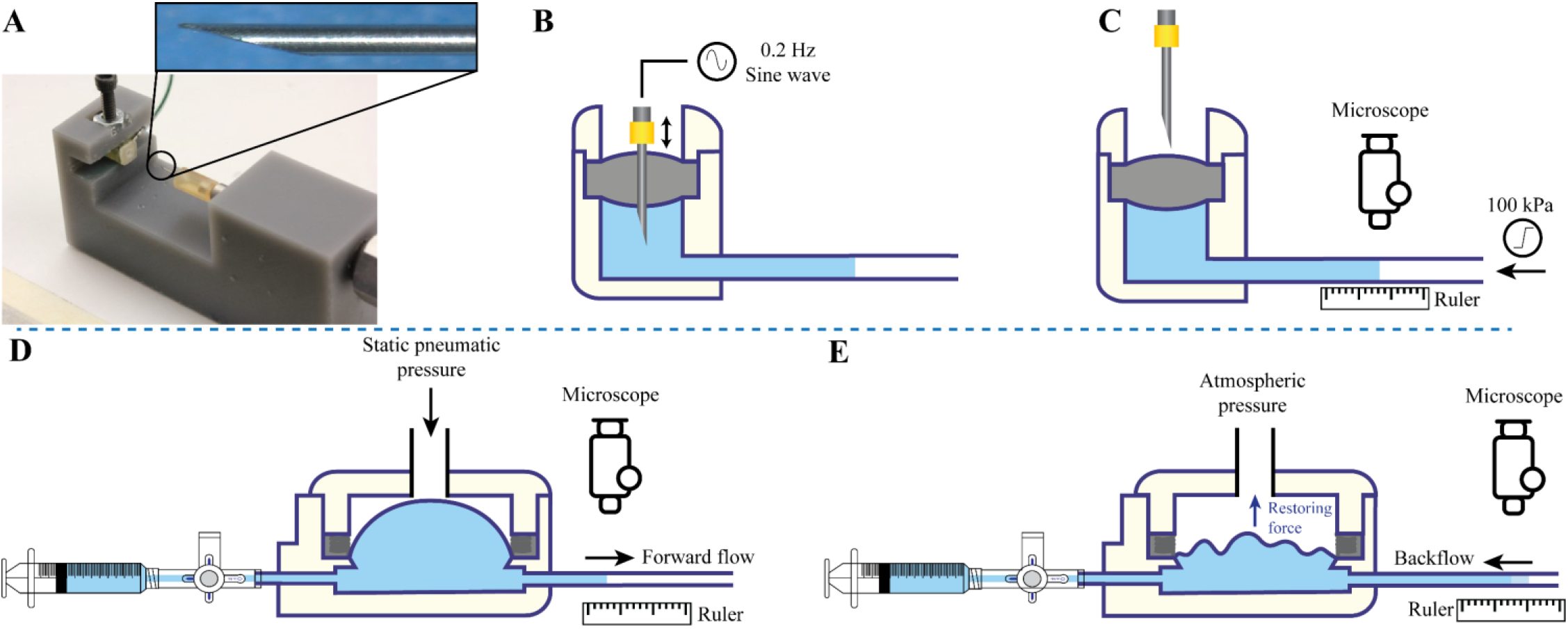
Benchtop characterization test setup for septum (Top) and cavity membrane (Bottom) characterization. Septum characterization: A) A custom-made 3D-printed pneumatic puncture device. Inset: a 30 Ga non-coring 12° beveled needle used for puncture test. B) The septum test samples were punctured in a single point following a logarithmic number of punctures with the puncture device fed by a 0.2 Hz sine wave signal. C) With the needle extracted, backpressure was gradually increased to 100 kPa while the fluid front displacement was quantified by visual observation under a microscope. Cavity characterization: D) Following filling of the microreservoir cavity and fluidically blocking the inlet, the fluid in the cavity membrane was pumped out via a static pneumatic pressure on the membrane to provide forward flow. E) The pneumatic pressure was released, and the backflow in the tubing due to the restoring force of the membrane was quantified under a microscope. Different tubing sizes were used for the three different cavity capacities (1, 10, 100 μL) to maintain a minimum resolution of 0.1% of the full microreservoir capacity.

Septum samples were prepared for puncture experiments with 27 and 30 Ga needles, where the septum cap incorporated the compression ring to compress the septum vertically and laterally for enhanced self-healing. To investigate the effect of the compression ring on the self-healing characteristics of the septum, one batch of samples (N=4) were fabricated without the septum cap. Instead, they were fixed and sealed in the refill port using cyanoacrylate, causing no compression to the septum. 30 Ga needles were used for this experiment, and the results were compared with the samples with the septum cap incorporating compression ring, punctured with 27 and 30 Ga needles (N=4 for each).

To test septum function, the space beneath the septum and 5 cm of the outlet tubing were filled with dyed DI water using the sharp (12°) non-coring needles, and the septum samples were fixed in the pneumatic puncture device. The outlet tubing was placed on a ruler to enable quantification of fluid motion for leak testing. A signal generator provided a 0.2 Hz sinusoidal drive to the pneumatic actuator to repeatedly puncture the septum. The number of repetitive punctures was increased logarithmically using a step size of 10^0.2^, with leak testing between each step. This leak testing involved gradual application of 100 kPa pneumatic back pressure at the outlet tubing for 1 min, with fluid displacement in the tubing observed/recorded with a digital microscope (USB-MICRO-250X, Plugable®, WA, USA) and analyzed with NIH-ImageJ, resulting in a resolution of 2.4 nL/min. Leakage was detected if a backward displacement of the fluid in the tubing was observed. If no leakage was observed the test continued by switching on the signal generator until the next step. The number of punctures before leakage was found for each case based on the last value with no leakage (Figure 4B).

**Figure 4.**
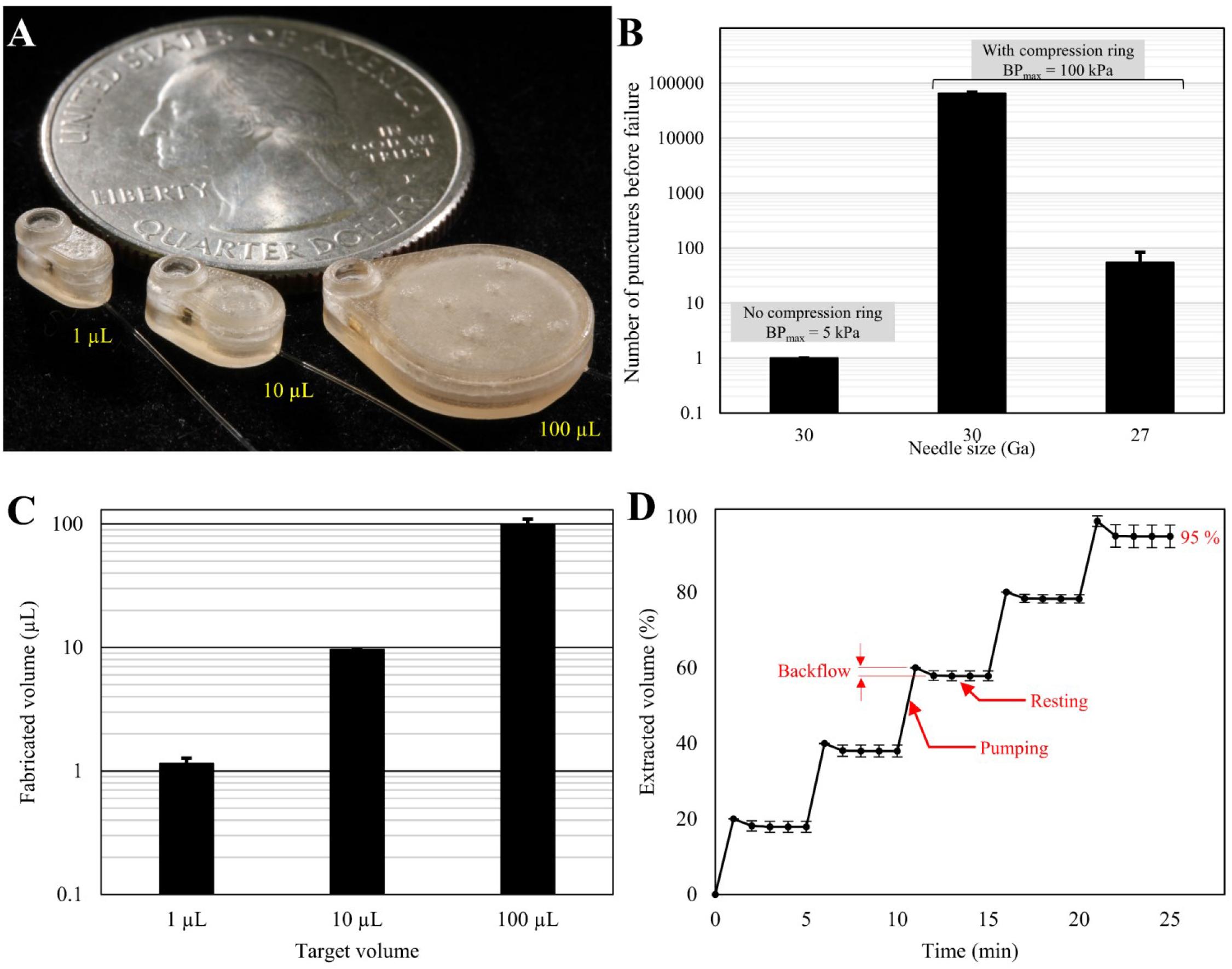
A) Photograph of completed microreservoirs with capacities of 1, 10, 100 μL, showing the scalability of the design and fabrication process. B) Septum characterization results show that the samples without the compression ring leaked at less than 5 kPa backpressure with just one puncture with a 30 Ga needle. Adding a compression ring to the septum cap increased the number of punctures before failure to ~65,000 at 100 kPa backpressure when puncturing with a 30 Ga needle. This design can provide no leakage up to 55 injections at 100 kPa backpressure when punctured by a 27 Ga needle, N=4, mean ± SD. C) Fabricated volumes were 1.15 ± 0.12, 9.63 ± 0.12, and 100.04 ± 9.43 for target volumes of 1, 10, and 100 μL, N=4, mean ± SD. D) The average results of the backflow due to restoring force for three different capacities of 1, 10, and 100 μL normalized by the total volume of each cavity. The results show that the overall backflow is not significant (2% average) and occurs quickly (< 2 min) after pneumatic pressure release, suggesting stable behavior of the cavity membrane in long-term. The last step of the experiment shows the total extraction percentage. N=27, mean ± SD.

#### 2.3.2. Cavity membrane test

Microreservoir cavity volume, backflow due to membrane restoring force, and total fluid extraction percentage for three different cavity sizes were characterized. The cavity samples were fabricated and tested separately, consisting of the cavity membrane (described in section 2.2), but with modifications to facilitate the test. The cavity inlet was directly connected to an inlet tubing, a three-way stopcock, and syringes for cavity filling, with an outlet tubing aligned over a ruler to quantify fluid movement due to pneumatic pressure applied to the region above the cavity membrane.

To fill the cavity, the pneumatic pressure was set to zero, and a syringe was connected to the inlet port and pulled to remove air from the cavity and pull the membrane down to its minimum volume. The syringe was replaced with a three-way stopcock connecting to the inlet tubing. Dyed DI water was injected to fill the inlet tube up to the entrance of the cavity via visual observation under a microscope. A small volume syringe was then connected via the stopcock to accurately quantify injected volume, with a 25 μL syringe (1702 LT SYR, Hamilton Co., NV, USA) used for the 1 and 10 μL cavities, and a 250 μL syringe (1725 TLL, Hamilton Co., NV, USA) used for the 100 μL cavity. With the outlet tubing open, the syringe was discharged until the fluid was observed to reach the microreservoir exit. The outlet tubing was clamped closed, and the syringe discharged to fill the cavity, confirmed by visual observation under a microscope. The three-way stopcock was switched to block flow on the inlet side, the outlet tubing clamp released, and the fluid volume in the outlet tubing quantified. The cavity volume was calculated by subtraction of the fluid volume in the outlet tubing from the injected fluid volume from the syringe.

The backflow due to restoring force of the cavity membrane was characterized for cavity fill volumes of ~ 80%, 60%, 40%, 20%, and 0%. Pneumatic pressure was applied above the cavity membrane for 1 min to induce forward fluid movement as shown in Figure 3D. After discharge of ~20% of the volume, the pneumatic pressure was released, and the fluid front displacement in the outlet tubing was observed under a microscope for 4 min (Figure 3E). For three different cavity sizes, three different outlet tubing sizes were used to have a minimum resolution of 0.1% of the full capacity using a 0.5-mm graded ruler. The backflow of the fluid front is recorded, and the experiment is repeated for the next 20% of the overall volume. The total fluid in the outlet tubing after the fifth step indicates the total extraction percentage. Three samples for each of three different cavity sizes of 1, 10, and 100 μL were tested with three replications (N=9 for each size, total N= 27).

### 2.4. In vivo experimental methods

The integrated microsystem (microreservoir + micropump) was tested *in vivo* to show its suitability of subcutaneous implantation, functionality for acute drug delivery to the mouse inner ear, and capability for long-term implantation. The drug delivery test was performed following a protocol developed by our laboratory [68], for the administration of sodium salicylate, leading to temporary hearing loss – reversible shifts of otoacoustic emissions thresholds and levels. Salicylate can act as a competitive antagonist at the anion-binding site of prestin [69], which causes reversible disruption of outer hair cell motility. Specifically, this disruption of prestin causes a reduction in distortion product otoacoustic emission (DPOAE) amplitudes and thresholds. Therefore, salicylate was delivered to the round window membrane (RWM) and auditory function was evaluated using DPOAE measurements. To evaluate long-term biocompatibility, the microreservoir was implanted for six months with frozen sections of surrounding tissue analyzed histologically for signs of inflammation.

#### 2.4.1. Subcutaneous implantation and biocompatibility

The anesthetic used was ketamine (120mg/kg body weight), xylazine (10 mg/kg body weight) in combination with topical application of 4% lidocaine for analgesia for the implant surgery. Specifically, a young adult CBA/CaJ mouse was positioned on a servo-controlled heating pad maintaining aseptic conditions, and when the proper plane of anesthetic was achieved (toe pinch, heart rate, respiratory rate), the upper back was then shaved and cleaned, and the microsystem was inserted subcutaneously in the center of the upper back via a small incision. Supplementary doses at one-third of the initial dose were administered as needed to maintain the proper levels of general anesthesia. Following insertion ventral to the dermis, Medical grade adhesive (Loctite 4206, Rocky Hill, CT) was used to secure the wound closure, along with several stitches to close the incision over the microsystem.

Following data collection, the mouse was euthanatized (Euthasol®, 0.22 ml/kg, IP) and then perfused with 4% paraformaldehyde fixative transcardially. Next, tissue samples were dissected from around the microsystem and prepared for standard frozen sectioning followed by hematoxylin + eosin (H&E) tissue staining to visualize cellular locations and structure around the microsystem. All animal procedures were approved by the University of South Florida Institutional Animal Care and Use Committee and were performed using the National Institutes of Health and veterinary standards of care.

#### 2.4.2. Inner ear drug delivery

Two young adult (2-4 months of age) CBA/CaJ mice bred and raised in-house were used for this study. After subcutaneous implantation of the microsystems in the center of the upper back (section 2.4.1), a bullaostomy surgery was performed to prepare a site for infusion of the salicylate into the middle ear cavity. A mixture of ketamine (120 mg/kg body weight) and xylazine (10 mg/kg body weight) injected via the intraperitoneal route to deeply anesthetize the mice for the bullaostomy surgery. The left ventral surface of the neck was then shaved and cleaned. Surgery was performed on the left (ipsilateral) ear following procedures described by Borkholder et al. [68]. During infusions, each mouse was immobilized using anesthesia with supplementary doses at one-third of the initial dose administered as needed to maintain the proper levels of general anesthesia.

Auditory function was assessed via automated DPOAE threshold measurements at F2 frequencies of 8.9 kHz, 13.5 kHz, 17.9 kHz, 24.6 kHz, 35.8 kHz, and 51.4 kHz. Measurements performed before the surgery were used for a baseline to compare subsequent DPOAE threshold shifts. Further details of auditory function assessment are presented in our previous work [62,70].

The salicylate solution consisted of NaCl (120 mM), KCl (3.5 mM), CaCl2 (1.5 mM), glucose (5.5 mM), HEPES buffer (4-(2-hydroxyethyl)-1-piperazineethanesulfonic acid, 20 mM), and sodium salicylate (50 mM). The pH was adjusted to 7.5 using NaOH. All solutions were prepared on the day of the experiment using sterile double-distilled water. The salicylate was loaded into a 25 μL sterilized syringe (1702 LT SYR, Hamilton Co., NV, USA) with a needle (33 Ga, 7747-01, point style 4, 12°, 1”, Hamilton Co., NV, USA) and was de-bubbled. The microreservoir was filled by injection through refill port using the needle and the salicylate was pushed through the tubing using positive syringe pressure until it was 1 mm from the microtubing tip.

The infusion of the salicylate was enabled by a novel biocompatible, implantable, scalable, and wirelessly controlled peristaltic micropump recently developed at our laboratory [62]. The mechanical components of the micropump were fabricated using direct-write 3D-printing technology on back of a printed circuit board assembly (PCBA), around the catheter microtubing outlet of the microreservoir to provide a biocompatible leak-free flow path while avoiding complicated microfluidic interconnects. The peristalsis is enabled by phase-change actuation of three chambers made adjacent to the microtubing. The actuation, control, and Bluetooth communication electronics were fabricated on the front side of the PCBA, effectively integrating mechanical and electronic components in a thin structure. The micropump operation is completely controlled through Bluetooth communication using a custom-made Android application. The micropump *in vitro* characterization results showed by tuning the actuation frequency of the micropump, nanoliter resolution flow rate across the range of 10-100 nL/min is generated, in the presence of 10× physiological inner ear backpressures and ±3 °C fluctuation in ambient temperature. It was also shown that the micropump could provide 50 nL/min for 20 min with 0.4% error in overall delivered drug volume, which was shown to temporarily impact the DPOAE threshold shift [62]. The integrated micropump operation was optimized for working in the subcutaneous temperature of mice (~33 °C) in a liquid environment (glycerin) with similar thermal characteristics to subcutaneous tissues to provide a flow rate of 50 nL/min for long-term performance.

## 3. Results and discussion

### 3.1. Benchtop experiments

The completed microreservoirs are shown in Figure 4A. The overall footprints of the microreservoirs are 5.8 × 3 (1 μL), 7.6 × 4.6 (10 μL), and 14.5 × 11.5 mm^2^ (100 μL) with a thickness of 3 mm for all. *In vitro* experiments were performed to optimize the septum design and thickness for thousands of refilling injections without leakage, and quantify backflow due to restoring force after extraction of liquid. Further, the total volume of the cavities and total extraction percentage of them were assessed. The cavity membrane experiments were performed on all three capacities of 1, 10, and 100 μL to demonstrate the scalability of the design.

#### 3.1.1. Septum characterization

The results (Figure 4B) indicated that the samples without the compression ring could not withstand one puncture and leaked at backpressures as low as 5 kPa. In contrast, the samples with the compression ring withstood up to ~65,000 punctures without any leakage at 100 kPa backpressure using the 30 Ga needle. However, using the 27 Ga needle reduced the number of punctures without leakage to 55. Comparison between the results of 30 Ga needle puncture with and without the compression ring indicates its significance in increasing the septum lifetime. Further, the results of 27 Ga punctures demonstrate the high endurance of this septum in taking up to 55 punctures, which is sufficient for many therapeutic development and clinical applications (e.g., [43,71]). Use of a 33 Ga needle (210 μm OD) dramatically enhances the robustness of this ultrathin septum to levels that far exceed any practical use case. This smaller gauge needle was successfully used in our *in vivo* experiments (section 3.2.1) with multiple successful subcutaneous injections in mice. All septum puncture experiments were performed on 1-mm thick septa. Testing on 0.5-mm thick septa showed poor functionality at low backpressures even with the compression ring. This could have been due to fabrication limitations for controlling the thickness of the compression ring and the buckling of the septum due to the lateral force of the compression. This novel septum design can be employed in any refillable system, independent of the cavity membrane design.

#### 3.1.2. Cavity membrane characterization

The results indicated that the overall capacities of the cavities were 1.15 ± 0.12, 9.63 ± 0.12, and 100.04 ± 9.43 (mean ± SD) for target volumes of 1, 10, and 100 μL, see Figure 4C. Figure 4D shows the average of extracted volume during pumping and resting for three different nominal capacities of 1, 10, and 100 μL, normalized by the total volume of each cavity. Each data point is an average of results from the three reservoir sizes, each with three samples and three replicates (N = 27). The results show that for all steps the backflow was not significant compared to the overall size (2% average). The backflow at each step occurs relatively quickly after deactivation of pumping (< 2 min for all cases), ensuring stable behavior of the cavity membrane for long-term applications. Further, no significant difference in the backflow was observed across different extraction percentage, except for the last step, which defines the total extraction percentage of the cavities. The average total extraction percentage was 95%.

Samples were also created with membranes twice the thickness of the original for each microreservoir size. Experiments (N=27) showed significant more backflow due to restoring force (3.8% average). This significant difference (p<0.001) is directly related to the thickness of the membranes. As the membrane becomes thicker, its resistance to deformation increases, which results in a higher restoring force and greater backflow when the pneumatic pressure is released. We hypothesize that the total extraction percentage is a function of the discrepancy between the surface area of the membrane and the substrate. If the membrane surface area is larger than the substrate, when fully pushed down by the pneumatic pressure the membrane wrinkles on the substrate and resists against further pressure. When the pneumatic pressure is released these areas induce restoring force and backflow. On the other hand, if the membrane surface area is smaller than the substrate, there is space between the membrane and the substrate that cannot be swept by the membrane unless with further pressure, which restores when the pressure is released and causes backflow. The fluid loss due to evaporation were not explored here because the *in situ* application of the device is subcutaneously implanted, which is a moisture-saturated environment.

### 3.2. In vivo experiments

The completed integrated microsystem is depicted in Figure 5A. The overall microsystem size measures 19 × 13 × 3 mm^3^ (L × W × H) enabling subcutaneous implantation in a mouse due to its planar form factor (Figure 5B). To evaluate the functionality of the microsystem, the outlet microtubing of the microsystem was implanted at the RWM niche to deliver salicylate at 50 nL/min for 20 min, replicating a previous drug delivery system utilizing a syringe pump [68]. Auditory function was measured during the infusion using the DPOAE method. Finally, the capability of the microsystem for long-term implantation was tested via standard frozen sectioning analysis of the tissue around it.

**Figure 5.**
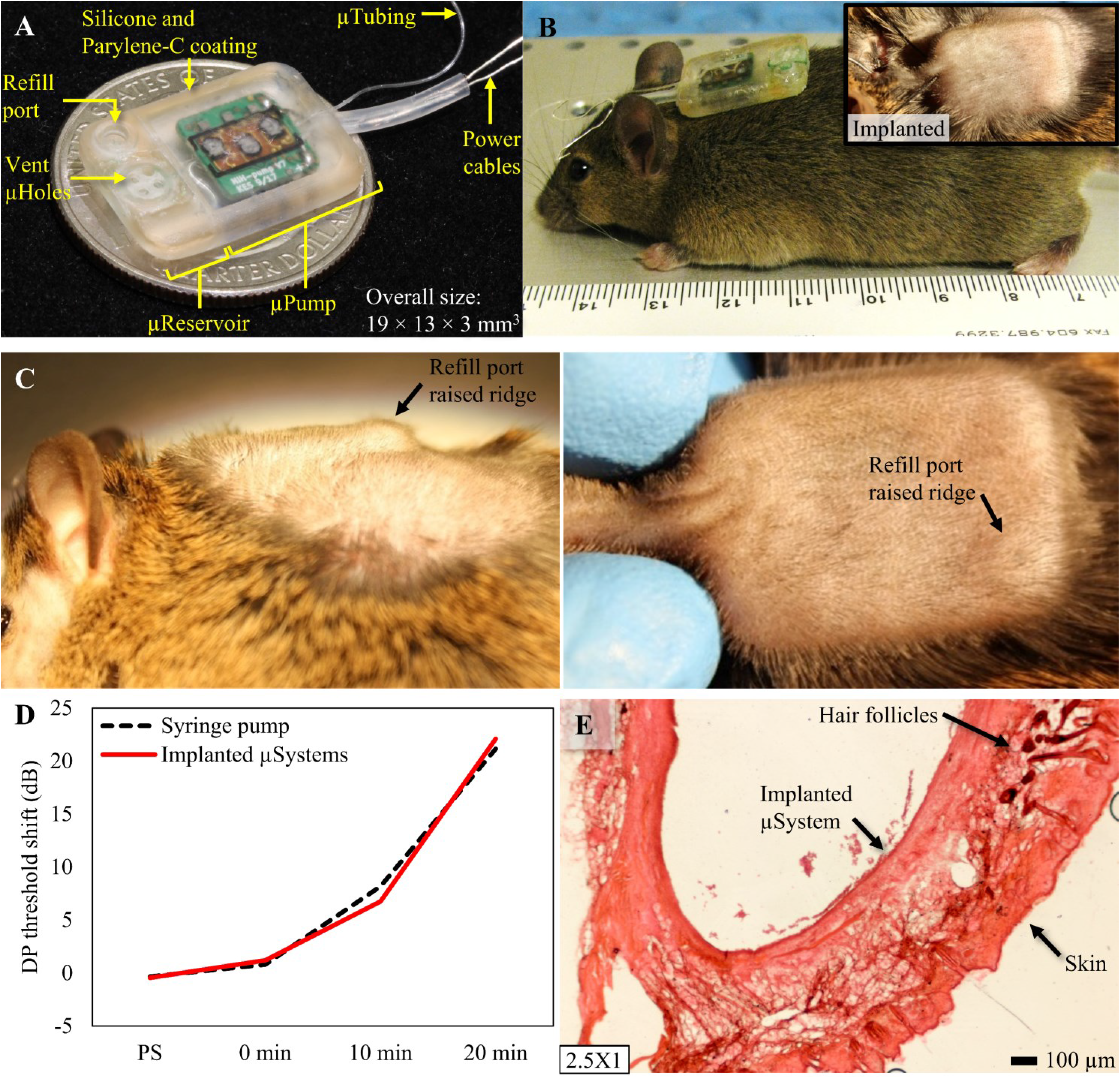
A) Photograph of the microreservoir integrated with a micropump for inner ear drug delivery that was recently reported [60]. The microsystem is optimized for subcutaneous implantation in mice with an overall thickness of 3 mm. B) Photograph of a microsystem before implantation demonstrates thinness of the microsystem, which makes it optimized for subcutaneous implantation in mice. Inset: An image of the microsystem after implantation via insertion of the microsystem in a small incision made in the center of the upper back of a mouse. C) Side view and top view of a microsystem implanted showing the visibility of the refilling port. D) Results of the mean threshold shift of the most basal region (F2 = 51.4 kHz) for implanted microsystems and the syringe pump show perfect consistency, demonstrating successful performance of the microsystem (N=6 for syringe pumps, N=2 for microsystems). E) A typical 60 μm-thick section of the skin surrounding the microsystem shows no sign of any active inflammatory cells. A fibrotic layer (pink) fully encapsulated the microsystem.

#### 3.2.1. Inner ear drug delivery

The accessibility of the refill port of the microreservoir was explored by palpation and observation of the raised ridge of the refill port when the shaved skin of the mouse is stretched (Figure 5C). Subcutaneous injections through the mouse skin were successfully performed by a 33 Ga (210 μm OD) non-coring needle (7747-01, point style 4, 12°, 1”, Hamilton Co., NV, USA), demonstrating *in vivo* validity of the larger 27 and 30 Ga syringe needles used for the septum characterization test.

Drug delivery experiments were performed using the integrated microsystem to show the performance of the system while implanted (two microsystems in two mice). Acute deliveries were run to deliver salicylate at the RWM niche at 50 nL/min for 20 min, while the DPOAE threshold shifts were recorded during the infusion, to replicate a previous experiment that was performed using a syringe pump [68]. The Android application enabled readily control of the device for staring/stopping the infusion. The results of the mean threshold shift of the most basal region (F2 = 51.4 kHz) for microsystems and the syringe pump are presented in Figure 5D. A baseline measurement was acquired approximately 10–15 min before the cannula placement surgery. The results compare the changes in DPOAE thresholds to baseline at four time-points: post-surgery (PS) in which DPOAE values were acquired immediately before the micropump was turned on, 0 min, the time at which the micropump was turned on, and 10 and 20 min after the onset of salicylate perfusion. There were mean shifts of 6.8 and 22.1 dB after 10 min and 20 min infusion for the implanted microsystems, consistent with the syringe pump results, suggesting the successful performance of the implanted microsystem.

#### 3.2.2. Long-term biocompatibility

The mouse recovered well from the surgery, and over the six-month survival period, was in good health. Post-implantation, the mouse was monitored daily and the results indicated that the overall health was excellent. This included no signs of fever or infection, no significant swelling, no redness or lumps around the micropump location or incision, nor presence of discolored fluid, and no weight loss. Cage behaviors, including feeding, grooming, and drinking were normal. The mouse was photographed on a weekly basis during the six months period.

The histological processing of the tissue surrounding the microsystem revealed a significant fibrotic layer around it, along with some ingrown hair follicles associated with the incision site. Otherwise, there were no cellular indications of infections or additional inflammatory responses, or abnormal cellular structures or responses. A typical section of the skin surrounding the microsystem is shown in Figure 5E. The skin surface is on the bottom, showing the normal dermal layers, with no sign of any active, inflammatory cells. The microsystem was in the cavity on the upper surface; here a fibrotic layer is seen (pink) which fully encapsulated the microsystem. This was the only abnormal features of the tissue surrounding the microsystem after six months survival, other than the dark hair follicles (upper right of the figure) associated with the incision site.

## 4. Conclusion

We developed a stand-alone refillable, scalable microreservoir designed for integration to an active pumping mechanism, optimized for subcutaneous implanted microsystems. The microreservoir structure has two main components: the refill port for refilling with subcutaneous injection through a septum, and the cavity area to store the drug for long-term applications. The microreservoir was built upon a 3D-printed biocompatible structure, providing mechanical protection and strength for handling, a stiff baseplate for the needle at the bottom of refill port, and a baseplate for the cavity area. Further, 3D-printing technology enables scalability of the design, supporting translational opportunities. The cavity membrane was fabricated with parylene using PEG as the sacrificial layer, over the parylene-coated substrate. The septum was fabricated using long-term biocompatible silicone rubber and coated with parylene. The biocompatibility of the system was strengthened by ensuring that all the materials are biocompatible, even the sacrificial layer.

The durability of the septum was significantly improved by adding a compression ring to the septum cap to induce internal compression in the septum. *In vitro* characterization results demonstrate no leakage after up to ~65,000 injections with a 30 Ga needle at up to 100 kPa backpressure, which is significantly higher than the maximum backpressure in humans and mice. The suitability of the needle size was verified by *in vivo* experiments which showed that needles as small as 33 Ga can be used for subcutaneous refills without breakage. This novel septum design can be used in different refillable microreservoirs. Scalability of the design and fabrication process was demonstrated by the fabrication of three different cavity sizes (1, 10, 100 μL) while keeping the overall thickness 3 mm. *In vivo* experiments demonstrated the suitability of this thickness for long-term (6 months) subcutaneous implantation in mice. *In vitro* characterization results show that the membrane causes an average backflow as low as 2% of the overall volume of the cavities. This feature permits integration of the microreservoir to normally-open micropumps for applications with low/zero backpressure (e.g., RWM in the inner ear). Further, when integrated to normally-closed micropumps, this design eliminates the long-term backpressure on check valves and improves pumping efficiency.

The integrability of the microreservoir was demonstrated by building a microsystem consisted of the microreservoir and a recently developed micropump, designed and optimized for inner ear drug delivery. Two microsystems were implanted in the center of the upper back of two mice, and the outlet microtubing was implanted at the RWM niche. The drug delivery was performed following a protocol previously developed in our laboratory for infusion of salicylate using a syringe pump, while the auditory function was assessed using the DPOAE method. The *in vivo* results show that the mean value of changes in the DP threshold shift for the implanted microsystems matches the shifts caused by infusion with a syringe pump, demonstrating the suitability of the microreservoir to be integrated to a pumping mechanism for subcutaneously implanted active drug delivery systems. Further, long-term biocompatibility of an implanted microsystem in a mouse was tested, showing very favorable signs of biocompatibility for long-term implantation, along with the development of the fibrotic layer surrounding the microsystem.

The presented microreservoir was shown to be refillable for thousands of injections with minimal backflow due to the discharging of the fluid when the pumping mechanism is off. Scalability of the design and fabrication process was demonstrated by the fabrication of three different cavity sizes (1, 10 100 μL), indicating translational opportunities. It was also shown that this microreservoir could be integrated with a pumping mechanism that either sucks the fluid from the outlet tubing (the integrated microsystem) or pushes the fluid directly from the top surface of membrane using pneumatic pressure (the cavity membrane test). Future work will incorporate a pumping mechanism from the membrane top surface for lab-on-chip or drug delivery applications.

## Acknowledgments

This work was supported by the National Institute on Deafness and Other Communication Disorders of the National Institutes of Health [grant number R01 DC014568].

## References

[1] C.L. Stevenson, J.T. Santini, R. Langer, Reservoir-based drug delivery systems utilizing microtechnology, Adv. Drug Deliv. Rev. 64 (2012) 1590–1602. doi:10.1016/j.addr.2012.02.005.

[2] J.H. Sakamoto, A.L. van de Ven, B. Godin, E. Blanco, R.E. Serda, A. Grattoni, A. Ziemys, A. Bouamrani, T. Hu, S.I. Ranganathan, E. De Rosa, J.O. Martinez, C.A. Smid, R.M. Buchanan, S.Y. Lee, S. Srinivasan, M. Landry, A. Meyn, E. Tasciotti, X. Liu, P. Decuzzi, M. Ferrari, Enabling individualized therapy through nanotechnology, Pharmacol. Res. 62 (2010) 57–89. doi:10.1016/j.phrs.2009.12.011.

[3] R.A.M. Receveur, F.W. Lindemans, N.F. De Rooij, Microsystem technologies for implantable applications, J. Micromechanics Microengineering. 17 (2007) R50–R80. doi:10.1088/0960-1317/17/5/R02.

[4] A.C.R. Grayson, R.S. Shawgo, A.M. Johnson, N.T. Flynn, Y. Li, M.J. Cima, R. Langer, A BioMEMS review: MEMS technology for physiologically integrated devices, in: Proc. IEEE, 2004: pp. 6–21. doi:10.1109/JPROC.2003.820534.

[5] M. Staples, K. Daniel, M.J. Cima, R. Langer, Application of micro- and nano-electromechanical devices to drug delivery, Pharm. Res. 23 (2006) 847–863. doi:10.1007/s11095-006-9906-4.

[6] A. Nisar, N. Afzulpurkar, B. Mahaisavariya, A. Tuantranont, MEMS-based micropumps in drug delivery and biomedical applications, Sensors Actuators, B Chem. 130 (2008) 917–942. doi:10.1016/j.snb.2007.10.064.

[7] Thomson Physicians’ Desk Reference, 63rd ed., Thomson Publishing, Montvale, NJ, 2009.

[8] Y.E. Choonara, V. Pillay, M.P. Danckwerts, T.R. Carmichael, L.C. Du Toit, A review of implantable intravitreal drug delivery technologies for the treatment of posterior segment eye diseases, J. Pharm. Sci. (2010). doi:10.1002/jps.21987.

[9] G.E. Sanborn, R. Anand, R.E. Torti, S.D. Nightingale, S.X. Cal, B. Yates, P. Ashton, T. Smith, Sustained-Release Ganciclovir Therapy for Treatment of Cytomegalovirus Retinitis: Use of an Intravitreal Device, Arch. Ophthalmol. (1992). doi:10.1001/archopht.1992.01080140044023.

[10] T.J. Smith, P.A. Pearson, D.L. Blandford, J.D. Brown, K.A. Goins, J.L. Hollins, E.T. Schmeisser, P. Glavinos, L.B. Baldwin, P. Ashton, Intravitreal Sustained-Release Ganciclovir, Arch. Ophthalmol. (1992). doi:10.1001/archopht.1992.01080140111037.

[11] P. Mruthyunjaya, D. Khalatbari, P. Yang, S. Stinnett, R. Tano, P. Ashton, H. Guo, M. Nazzaro, G.J. Jaffe, Efficacy of low-release-rate fluocinolone acetonide intravitreal implants to treat experimental uveitis, Arch. Ophthalmol. (2006). doi:10.1001/archopht.124.7.1012.

[12] D.G. Callanan, G.J. Jaffe, D.F. Martin, P.A. Pearson, T.L. Comstock, Treatment of posterior uveitis with a fluocinolone acetonide implant: Three-year clinical trial results, Arch. Ophthalmol. (2008). doi:10.1001/archopht.126.9.1191.

[13] B.D. Kuppermann, Implant delivery of corticosteroids and other pharmacologic agents, Retina. (2006) 10–11.

[14] A.C. Richards Grayson, I.S. Choi, B.M. Tyler, P.P. Wang, H. Brem, M.J. Cima, R. Langer, Multi-pulse drug delivery from a resorbable polymeric microchip device, Nat. Mater. (2003). doi:10.1038/nmat998.

[15] G.Y. Kim, B.M. Tyler, M.M. Tupper, J.M. Karp, R.S. Langer, H. Brem, M.J. Cima, Resorbable polymer microchips releasing BCNU inhibit tumor growth in the rat 9L flank model, J. Control. Release. (2007). doi:10.1016/j.jconrel.2007.08.003.

[16] A.C.R. Grayson, M.J. Cima, R. Langer, Molecular release from a polymeric microreservoir device: Influence of chemistry, polymer swelling, and loading on device performance, J. Biomed. Mater. Res. – Part A. (2004). doi:10.1002/jbm.a.30019.

[17] http://www.debiotech.com/., (n.d.). http://www.debiotech.com/.

[18] L.-D. Piveteau, H. Hofmann, F. Neftel, Anisotropic nanoporous coatings for medical implants, (2009).

[19] D. Capodanno, F. Dipasqua, C. Tamburino, Novel drug-eluting stents in the treatment of de novo coronary lesions, Vasc. Health Risk Manag. (2011). doi:10.2147/VHRM.S11444.

[20] E. Grube, J. Schofer, K.E. Hauptmann, D. Nickenig, N. Curzen, D.J. Allocco, K.D. Dawkins, A novel paclitaxel-eluting stent with an ultrathin abluminal biodegradable polymer: 9-month outcomes with the jactax hd stent, JACC Cardiovasc. Interv. (2010). doi:10.1016/j.jcin.2009.12.015.

[21] P.J.T. A.N. Paisley, Recent advances in the treatment of acromegaly, Recent Adv. Investig. Drugs. 15 (2006) 251–256.

[22] F. Martin, R. Walczak, A. Boiarski, M. Cohen, T. West, C. Cosentino, M. Ferrari, Tailoring width of microfabricated nanochannels to solute size can be used to control diffusion kinetics, J. Control. Release. (2005). doi:10.1016/j.jconrel.2004.09.024.

[23] P. Gardner, Use of a nanopore membrane in novel a drug delivery device, Futur. Drug Deliv. (2006) 59–60.

[24] C.L. Stevenson, F. Theeuwes, J.C. Wright, Osmotic implantable delivery systems, Handb. Pharm. Control. Release Technol. (2000).

[25] J.C. Wright, S.T. Leonard, C.L. Stevenson, J.C. Beck, G. Chen, R.M. Jao, P.A. Johnson, J. Leonard, R.J. Skowronski, An in vivo/in vitro comparison with a leuprolide osmotic implant for the treatment of prostate cancer, J. Control. Release. 75 (2001) 1–10.

[26] C.L. Stevenson, Formulation of leuprolide at high concentration for delivery from a one-year duration implant, in: Protein Formul. Deliv. Second Ed., CRC Press, 2007: pp. 171–194.

[27] D.M. Fisher, N. Kellett, R. Lenhardt, Pharmacokinetics of an implanted osmotic pump delivering sufentanil for the treatment of chronic pain, Anesthesiology. (2003). doi:10.1097/00000542-200310000-00028.

[28] www.intarcia.com., (n.d.). www.intarcia.com.

[29] T.A. B. Yang, C. Negulescu, R. D’vas, C. Eftimie, J. Carr, S. Lautenbach, K. Horwege, C. Mercer, D. Ford, Stability of ICTA 560 for continuous subcutaneous delivery of Exenatide at body temperature for 12 months, in: 2009 Diabetes Technol. Meet., Long Beach, CA, 2009.

[30] H. R.R., C. R., R. J., A. T., L. K., Comparing ITCA 650, continuous subcutaneous delivery of exenatide via DUROS(registered trademark) device, vs. twice daily exenatide injections in metformintreated type 2 diabetes, in: Diabetologia, 2010.

[31] D.F. B. Yang, T. Alessi, C. Rohloff, R. Mercer, C. Negulescu, S. Lautenbach, J. Gumucio, M. Guo, E. Weeks, J. Carr, Continuous delivery of proteins and peptides at consistent rates for at least 3 months form the DUROS® device,, Poster T3150, in: AAPS 2008 Meet., Atlanta Georgia, n.d.

[32] C.M. Rohloff, T.R. Alessi, B. Yang, J. Dahms, J.P. Carr, S.D. Lautenbach, DUROS®technology delivers peptides and proteins at consistent rate continuously for 3 to 12 months, J. Diabetes Sci. Technol. (2008). doi:10.1177/193229680800200316.

[33] C. Abe, T. Tashiro, K. Tanaka, R. Ogihara, H. Morita, A novel type of implantable and programmable infusion pump for small laboratory animals, J. Pharmacol. Toxicol. Methods. 59 (2009) 7–12. doi:10.1016/j.vascn.2008.09.002.

[34] T. Tan, S.W. Watts, R.P. Davis, Drug delivery: Enabling technology for drug discovery and development. iPRECIO® Micro Infusion Pump: programmable, refillable, and implantable, Front. Pharmacol. JUL (2011) 44. doi:10.3389/fphar.2011.00044.

[35] R. Sheybani, A. Cobo, E. Meng, Wireless programmable electrochemical drug delivery micropump with fully integrated electrochemical dosing sensors, Biomed. Microdevices. 17 (2015) 1–13. doi:10.1007/s10544-015-9980-7.

[36] A.J. Chung, Y.S. Huh, D. Erickson, A robust, electrochemically driven microwell drug delivery system for controlled vasopressin release, Biomed. Microdevices. 11 (2009) 861–867. doi:10.1007/s10544-009-9303-y.

[37] N.M. Elman, H.L. Ho Duc, M.J. Cima, An implantable MEMS drug delivery device for rapid delivery in ambulatory emergency care, Biomed. Microdevices. 11 (2009) 625–631. doi:10.1007/s10544-008-9272-6.

[38] R. Lo, P.-Y. Li, S. Saati, R. Agrawal, M.S. Humayun, E. Meng, A refillable microfabricated drug delivery device for treatment of ocular diseases, Lab Chip. 8 (2008) 1027. doi:10.1039/b804690e.

[39] R. Lo, P.-Y. Li, S. Saati, R.N. Agrawal, M.S. Humayun, E. Meng, A passive MEMS drug delivery pump for treatment of ocular diseases., Biomed. Microdevices. 11 (2009) 959–70. doi:10.1007/s10544-009-9313-9.

[40] A. Cobo, R. Sheybani, H. Tu, E. Meng, A wireless implantable micropump for chronic drug infusion against cancer, Sensors Actuators A Phys. 239 (2016) 18–25. doi:10.1016/j.sna.2016.01.001.

[41] H. Gensler, R. Sheybani, P.Y. Li, R. Lo Mann, E. Meng, An implantable MEMS micropump system for drug delivery in small animals, Biomed. Microdevices. 14 (2012) 483–496. doi:10.1007/s10544-011-9625-4.

[42] Debiotech, (n.d.). https://www.debiotech.com (accessed October 14, 2018).

[43] E. Zini, I. Padrutt, K. Macha, A. Riederer, M. Pesaresi, T.A. Lutz, C.E. Reusch, Use of an implantable pump for controlled subcutaneous insulin delivery in healthy cats, Vet. J. 219 (2017) 60–64. doi:10.1016/j.tvjl.2016.12.006.

[44] F.N. Pirmoradi, J.K. Jackson, H.M. Burt, M. Chiao, On-demand controlled release of docetaxel from a battery-less MEMS drug delivery device, Lab Chip. 11 (2011) 2744–2752. doi:10.1039/c1lc20134d.

[45] Y.B. Gianchandani, S. Chiravuri, R.Y. Gianchandani, T. Li, A.T. Evans, Compact, power-efficient architectures using microvalves and microsensors, for intrathecal, insulin, and other drug delivery systems, Adv. Drug Deliv. Rev. 64 (2012) 1639–1649. doi:10.1016/j.addr.2012.05.002.

[46] D. Reynaerts, J. Peirs, H. Van Brussel, K.U. Leuven, A SMA-ACTUATED IMPLANTABLE SYSTEM FOR DELIVERY OF LIQUID DRUGS, (1996) 26–28.

[47] M. Wook, S. An, S.S. Yoon, A.L. Yarin, Advances in self-healing materials based on vascular networks with mechanical self-repair characteristics, 252 (2018) 21–37. doi:10.1016/j.cis.2017.12.010.

[48] R.F. Shepherd, A.A. Stokes, R.M.D. Nunes, G.M. Whitesides, Soft machines that are resistant to puncture and that self seal, Adv. Mater. 25 (2013) 6709–6713. doi:10.1002/adma.201303175.

[49] L.E.F. Circuits, P. Elast, Re-configu le fluid circuits, Science (80-.). (n.d.) 222–227.

[50] M.K. Chaudhury, G.M. Whitesides, Direct Measurement of Interfacial Interactions between Semispherical Lenses and Flat Sheets of Poly(dimethylsiloxane) and Their Chemical Derivatives, Langmuir. 7 (1991) 1013–1025. doi:10.1021/la00053a033.

[51] J.C. Andrews, S.C. Walker-Andrews, W.D. Ensminger, Long-term central venous access with a peripherally placed subcutaneous infusion port: initial results., Radiology. 176 (1990) 45–47. doi:10.1148/radiology.176.1.2353108.

[52] S. S., M. J., K. A., J. C., Improved methods for venous access: The Port-A-Cath, a totally implanted catheter system, J. Clin. Oncol. 4 (1986) 596–603. https://www.researchgate.net/profile/Stephen_Strum/publication/284291764_The_Port-a-cath_PAC_A_totally_implanted_catheter_system/links/56aa5b0008aef6e05df46165/The-Port-a-cath-PAC-A-totally-implanted-catheter-system.pdf (accessed April 1, 2019).

[53] iPRECIO^®^, (n.d.). http://www.iprecio.com.

[54] Prometra Pump | Flowonix, (n.d.). http://www.flowonix.com/healthcare-provider/products/prometra-pump.

[55] ithetis Implantable Drug Delivery Devices, (n.d.). http://www.ithetis.com.

[56] R. Lo, K. Kuwahara, P.Y. Li, R. Agrawal, M.S. Humayun, E. Meng, Ieee, A passive refillable Intraocular MEMS drug delivery device, Int. Conf. Microtechnologies Med. Biol. (2006) 74–77. doi:10.1109/MMB.2006.251494.

[57] P.Y. Li, J. Shih, R. Lo, S. Saati, R. Agrawal, M.S. Humayun, Y.C. Tai, E. Meng, An electrochemical intraocular drug delivery device, Sensors Actuators, A Phys. 143 (2008) 41–48. doi:10.1016/j.sna.2007.06.034.

[58] A.T. Evans, S. Chiravuri, Y.B. Gianchandani, A Multidrug Delivery System Using a Piezoelectrically Actuated Silicon Valve Manifold With Embedded Sensors, J. Microelectromechanical Syst. 20 (2011) 231–238. doi:10.1109/JMEMS.2010.2093566.

[59] F.L. Trull, B.A. Rich, More regulation of rodents, Science (80-.). (1999). doi:10.1126/science.284.5419.1463.

[60] E. Benedikz, E. Kloskowska, B. Winblad, The rat as an animal model of Alzheimer’s disease, J. Cell. Mol. Med. (2009). doi:10.1111/j.1582-4934.2009.00781.x.

[61] J. Urquhart, Rate-Controlled Delivery Systems in Drug and Hormone Research, Annu. Rev. Pharmacol. Toxicol. 24 (2002) 199–236. doi:10.1146/annurev.pharmtox.24.1.199.

[62] F. Forouzandeh, X. Zhu, A. Alfadhel, B. Ding, J.P. Walton, D. Cormier, R.D. Frisina, D.A. Borkholder, A nanoliter resolution implantable micropump for murine inner ear drug delivery, J. Control. Release. 298 (2019) 27–37. doi:10.1016/j.jconrel.2019.01.032.

[63] D.J. Laser, J.G. Santiago, A review of micropumps, J. Micromechanics Microengineering. 14 (2004) R35–R64. doi:10.1088/0960-1317/14/6/R01.

[64] R. Lo, E. Meng, Integrated and reusable in-plane microfluidic interconnects, Sensors Actuators, B Chem. 132 (2008) 531–539. doi:10.1016/j.snb.2007.11.024.

[65] H.M. Gensler, H.M. Gensler, A Wireless Implantable MEMS Micropump System for Site-specific Anti-cancer Drug Delivery by Copyright 2013, (2013).

[66] S. coating Systems, SCS Parylene properties, (n.d.). https://scscoatings.com.

[67] Hamilton reference guide – Syringes and Needles, (n.d.). www.hamiltoncompany.com.

[68] D.A. Borkholder, X. Zhu, R.D. Frisina, Round window membrane intracochlear drug delivery enhanced by induced advection, J. Control. Release. 174 (2014) 171–176. doi:10.1016/j.jconrel.2013.11.021.

[69] D. Oliver, D.Z.Z. He, N. Klöcker, J. Ludwig, U. Schulte, S. Waldegger, J.P. Ruppersberg, P. Dallos, B. Fakler, Intracellular anions as the voltage sensor of prestin, the outer hair cell motor protein, Science (80-.). 292 (2001) 2340–2343. doi:10.1126/science.1060939.

[70] R.D. Frisina, B. Ding, X. Zhu, J.P. Walton, Age-related hearing loss: Prevention of threshold declines, cell loss and apoptosis in spiral ganglion neurons, Aging (Albany. NY). 8 (2016) 2081–2099. doi:10.18632/aging.101045.

[71] R. Rauck, T. Deer, S. Rosen, G. Padda, J. Barsa, E. Dunbar, G. Dwarakanath, Accuracy and efficacy of intrathecal administration of morphine sulfate for treatment of intractable pain using the Prometra® Programmable Pump, Neuromodulation. 13 (2010) 102–107. doi:10.1111/j.1525-1403.2009.00257.x.

